# A novel class of sulfur-containing aminolipids widespread in marine roseobacters

**DOI:** 10.1101/2021.02.05.429882

**Authors:** Alastair F. Smith, Eleonora Silvano, Orsola Päuker, Richard Guillonneau, Mussa Quareshy, Andrew Murphy, Michaela A Mausz, Rachel Stirrup, Branko Rihtman, Maria Aguilo Ferretjans, Joost Brandsma, Jörn Petersen, David J Scanlan, Yin Chen

## Abstract

Marine roseobacter group bacteria are numerically abundant and ecologically important players in ocean ecosystems. These bacteria are capable of modifying their membrane lipid composition in response to environmental change. Remarkably, a variety of lipids are produced in these bacteria, including phosphorus-containing glycerophospholipids and several amino acid-containing aminolipids such as ornithine lipids and glutamine lipids. Here, we present the identification and characterization of a novel <u>s</u>ulfur-containing <u>a</u>mino<u>l</u>ipid (SAL) in roseobacters. Using high resolution accurate mass spectrometry, a SAL was found in the lipid extract of *Ruegeria pomeroyi* DSS-3 and *Phaeobacter inhibens* DSM 17395. Using comparative genomics, transposon mutagenesis and targeted gene knockout, we identified a gene encoding a putative lyso-lipid acyltransferase, designated *SalA*, which is essential for the biosynthesis of this SAL. Multiple sequence analysis and structural modelling suggest that SalA is a novel member of the lysophosphatidic acid acyltransferase (LPAAT) family, the prototype of which is the PlsC acyltransferase responsible for the biosynthesis of the phospholipid phosphatidic acid. SAL appears to play a key role in biofilm formation in roseobacters. *SalA* is widely distributed in *Tara* Oceans metagenomes and actively expressed in *Tara* Oceans metatranscriptomes. Our results raise the importance of sulfur-containing membrane aminolipids in marine bacteria.

## Introduction

Bacterial lipids are the key constituent segregating cellular components from the external environment. Bacterial lipids are highly diverse yet there is currently little understanding of the benefits that this diversity provides (Sohlenkamp and Geiger, 2016). Glycerophospholipids are by far the best studied lipids in bacteria and a key branch point for glycerophospholipid biosynthesis is phosphatidic acid (PA), from which a variety of lipids, including phosphatidylglycerol (PG), phosphatidylethanolamine (PE), phosphatidylcholine (PC), diacylglycerol (DAG) and triacylglycerol (TAG), can be made through either the cytidine diphosphate (CDP)-diacylglycerol (DAG) pathway or the Kennedy pathway (Parsons & Rock, 2013). PA biosynthesis in bacteria is carried out by a membrane-attached acyltransferase PlsC, the founding member of the large lysophosphatidic acid acyltransferase (LPAAT) family (Robertson et al., 2017; Rottig & Steinbuchel 2013).

Aside from phospholipids, the study of bacterial lipid diversity is currently hampered by a lack of knowledge of both the chemical structures of many of these lipids and the identity of genes involved in their synthesis. These have severely hindered our understanding of lipid diversity and their physiological function in bacteria. Once the chemical structure of a lipid is known, analytical strategies can then be devised to detect the lipid in both the natural environment and cell cultures (Moore *et al.*, 2016). This can also help to direct studies into the biosynthesis of the lipid, knowledge of which can provide a clearer idea of the likely distribution of the lipid amongst various bacterial classes. A group of poorly studied bacterial lipids are the aminolipids, of which only ornithine lipids have been detected in diverse cultured bacteria since the 1960s (Shively and Knoche, 1969). However, it was not until the genes involved in its biosynthesis were elucidated that it became clear how widespread the capacity to produce ornithine lipid really was (Gao *et al.*, 2004; Vences-Guzmán *et al.*, 2015). Similarly, Sebastian *et al.* (2016) found several uncharacterised aminolipids in marine heterotrophic bacteria one of which was recently determined as a glutamine-containing aminolipid, often found in the marine roseobacter group (Smith et al., 2019). Both ornithine and glutamine lipids play a key role in the adaptation of cosmopolitan marine bacteria (e.g. the marine SAR11 clade and the roseobacter group) to oligotrophic environments (Carini et al., 2015; Sebastian et al., 2016; Smith et al., 2019).

In this study, we report the characterisation and chemical structure of a novel sulfur-containing aminolipid using high resolution–accurate mass spectrometry from the marine roseobacter group. This newly identified lipid represents a novel class of sulfur-containing lipids with an aminosulfonate head group. Furthermore, we describe a novel acyltransferase enzyme (SalA), part of the LPAAT family, that is responsible for the biosynthesis of this sulfonolipid. This sulfonolipid appears widespread within the roseobacter group that are key players in marine biogeochemical cycles and important for biofilm formation. Furthermore, the *salA* gene is abundant and actively transcribed in marine surface microbial assemblages.

## Materials and methods

### Bacterial strains and cultivation

All marine bacteria used in this study were cultivated using either marine broth medium (BD Difco™ 2216), ½YTSS medium containing yeast extract (2 g/L), tryptone (1.25 g/L), and Sigma sea salts (20 g/L) or a defined marine ammonium mineral salts (MAMS) medium (Smith et al., 2019). The MAMS medium contained 30 g/L NaCl, 10 mM glucose, 1 mM K_2_HPO_4_, 0.75-7.5 mM NH_4_Cl, 10 mM HEPES buffer (pH 7.6), 1.36 mM CaCl_2_, 0.98 mM MgSO_4_, 7.2 μM FeCl_2_, 84 μM Na_2_MoO_4_,370 nM ZnCl_2_, 510 nM MnCl_2_, 97 nM H3BO_3_, 1.1 μM CoCl_2_, 12 nM CuCl_2_, 100 nM NiCl_2_, 30 nM thiamine, 160 nM nicotinic acid, 97 nM pyridoxine, 73 nM aminobenzoic acid, 53 nM riboflavin, 84 nM pantothenate, 4.1 nM biotin, 1.5 nM cyanocobalamin and 11 nM folic acid. All cultures were grown at 30°C aerobically in a shaker (150 r.p.m) unless stated otherwise.

### Intact polar lipid analysis

Lipid extraction from bacterial cultures was carried out using the modified Folch extraction protocol as described previously (Smith et al., 2019). Briefly 1 ml culture of OD_540_~1.0 was collected by centrifugation. Total lipids were then extracted using methanol-chloroform, dried under nitrogen gas and the pellet re-suspended in 1 mL solvent (95% (v/v) liquid chromatography-mass spectrometry (LC-MS) grade acetonitrile and 5% 10 mM ammonium acetate pH 9.2 in water). These lipids were then analysed by LC-MS using a Dionex 3400RS HPLC with a HILIC BEH amide XP column (2.5 μm, 3.0×150 mm, Waters) coupled with an amaZon SL ion trap MS (Bruker) via electrospray ionisation (ESI) in both positive (+ve) and negative (-ve) ionisation mode. Samples were run on a 15 min gradient from 95% (v/v) acetonitrile/5% (w/v) ammonium acetate (in water, 10 mM, pH 9.2) to 70% (v/v) acetonitrile/30% (w/v) ammonium acetate (in water, 10 mM, pH 9.2), followed by 5 min of isocratic run 70% acetonitrile/30% ammonium acetate with 10 minutes equilibration between samples. The flow rate was maintained at 150 μL min^-1^ and the column temperature at 30°C. The injection volume was 5 μL for each run; the ionization was done in both positive and negative mode. Drying conditions were the same for both modes (8 L min^-1^ drying gas at 300°C and nebulizing gas pressure of 15 psi). The end cap voltage was 4,500 V in positive mode and 3,500 V in negative mode, both with 500 V offset. Data analysis was carried out using the Bruker Compass software package. Unless state otherwise, base peak chromatographs were presented with *m/z* range from 400 -1000.

High resolution MS identification and fragmentation was carried out using either a quadrupole-time-of-flight MS (Q-TOF, Waters Synapt G2-Si) or an Orbitrap Fusion (Thermo Fisher Scientific) by direct infusion and collision induced dissociation (CID). For the Orbitrap Fusion, the resolution was set at 120K with CID for MS^n^. A TriVersa Nanomate nanospray source (Advion, NY) was used and the flow rate was at 300 nl min^-1^. The voltage was set at 1.4 kV and the gas pressure was 0.3 psi. Sheath and sweep gas were set to zero and the cone voltage was 2100 V and the mass range was from 50-1000 Da. MS data were analyzed using Xcalibur (Thermo Fisher Scientific). For Q-TOF, samples were injected through a Universal NanoFlow Sprayer (Waters) by direct infusion at 200-300 nl min^-1^ and the cone voltage was 30 V in negative mode ESI. Mass range was set from 50-1000 Da and data analyses ware carried out in MassLynx (Waters). The most abundant peak in the negative ion spectrum corresponding to the SAL lipid (*m/z* 656.6) was selected for MS^n^ fragmentation. Spectra were obtained in profile mode and smoothed using a moving mean. Background correction using a linear baseline was applied with a 40% noise cut-off. For accurate mass determination, the centroid of each peak was used. The peak corresponding to C17H33COO- (*m/z* 281.2480, an 18:1 fatty acid carboxylate anion) was used as a lock mass. Calculation of candidate elemental formulae from the accurate mass considered formulae containing C0-100, H0-100, N0-100, S0-4, P0-1. A conservative mass error of 100 ppm was assumed.

### Marker-exchange mutagenesis

Marker-exchange mutagenesis was carried out as described previously using a suicide vector *pK18mobsacB* (Smith et al., 2019). Briefly, DNA fragments corresponding to an upstream element and a downstream element that flank the target gene were amplified by PCR using high-fidelity *Phusion* DNA polymerase. A Gm-resistance cassette was amplified from plasmid p34S-Gm (Dennis & Zylstra 1998; Smith et al., 2019). These fragments, together with the linearized *pK18mobsacB* vector were then assembled through Gibson cloning and transformed into competent *Escherichia coli* DH5α cells. The engineered suicide vector was then extracted from *E. coli* DH5α and transformed into the conjugation donor strain *E. coli* S17.1 *λpir* before conjugating into *Ruegeria pomeroyi* DSS-3 as described previously (Smith et al., 2019). Transconjugants were then selected on defined MAMS medium containing gentamycin (Gm, 10 μg/ml). All mutants were confirmed by PCR using the confirmation primers (**Supplementary Table 1**) and subsequent Sanger sequencing.

### Transposon library of *Phaeobacter inhibens* DSM 17395

A library of 5,500 transposon mutants of *Phaeobacter inhibens* DSM 17395, which was established at the DSMZ, served as a basis to identify genes involved in the biosynthesis of the novel SAL lipid. Transposon mutagenesis was performed with the EZ-Tn5 <R6Kγori/KAN-2>Tnp Transposome kit (Epicentre, Illumina, CA, USA) and the insertion site of all mutants was determined via arbitrary PCR (Segev et al. 2016). Transposon mutants were streaked out three times to eliminate attached wild type cells. The absence of wild type cells and the presence of the 65 kb plasmid were validated as described previously (Segev et al. 2016; Frank et al., 2015; Michael et al., 2016). The transposon integration site of each mutant was also confirmed via sequencing of the amplification PCR product, and stable maintenance of all three extrachromosomal elements was validated via diagnostic PCR (Trautwein et al., 2016).

The transposon mutant #1036 of *P. inhibens* DSM 17395 (PGA1_c01210) unable to produce the SAL lipid was complemented using the *salA* homologue of *Ruegeria pomeroyi* DSS-3 (locus tag SPO0716) and *P. inhibens* DSM 17395 (locus tag PGA1_c01210). Complementation was carried out by PCR amplification of the *salA* homologues together with a constitutive promoter (~250 bp upstream of the *aacC1* gene from plasmid p34S-GM, Dennis & Zylstra 1998), which was then cloned into the broad host range vector pBBR1MCS and transformed into the *salA* mutant of *P. inhibens* DSM 17395 by conjugation as described previously (Sebastian et al., 2016; Smith et al., 2019). The complemented mutants were cultivated using marine broth medium and cells were harvested for lipidomics analysis as described above.

### Biofilm assays

To grow biofilms of *Phaeobacter inhibens* DSM 17395 and the *salA* mutant, post-exponential grown bacterial cells were washed and diluted in fresh marine broth medium and inoculated at an OD_590 nm_ of 0.2 into 24 well plates (Corning Incorporated Costar®, New York, NY, United States) containing a sterilized glass coverslip into each well. At each time point (3, 24, and 48 h), biofilms were washed to remove non-adherent bacteria and fixed using formalin 3.5% (v/v) for 20 min. Bacteria were stained using DAPI (5 μg/mL, Sigma-Aldrich, Darmstadt, Germany) and coverslips were mounted with a drop of Mowiol antifade before observation using confocal laser scanning microscopy (CLSM) (Zeiss LSM 880, Göttingen, Germany). The biovolume and the average thickness of the biofilms were determined using COMSTAT software developed in MATLAB R2017a (MathWorks, Natick, MA, United States) as described previously (Heydorn et al., 2000; Guillonneau et al., 2018). To test for statistically significant differences between the wild-type strain and the *salA* mutant, a *t*-test was performed using SPSS 13.0 (IBM, Armonk, NY, United States).

A crystal violet biofilm assay was also performed which was adapted from Guillonneau et al. (2018). Bacterial biofilms were developed in 96-well microtiter plates (Greiner Bio-One, Kremsmünster, Austria) with bacteria in the post-exponential growth phase using marine broth medium. Cells were diluted to a final OD_590 nm_ = 0.1 into each well (n=4 for both the wild type and the *salA* mutant) and grown in static conditions at 30°C. At each time point (3 h, 24 h, 48 h and 72 h) samples were washed three times with fresh marine broth medium and dried for 30 min at 50°C. Biofilms were then stained for 15 min with 200 μL crystal violet 0.01% (w/v), rinsed three times with phosphate-buffered saline and dried for 10 min. Biofilm quantification was performed by releasing the stain from the biofilm using absolute ethanol for 10 min at 30°C with gentle shaking. The absorbance of the crystal violet in solution was measured at 595 nm. The final absorbance of each sample was calculated by subtracting the blank (*i.e.*, marine broth medium only treated with crystal violet, n=4).

### Bioinformatics analysis

Phylogenetic analysis of 16S rRNA genes from *Rhodobacteraceae* was carried out using the full length 16S rRNA gene retrieved from the Integrated Microbial Genomes (IMG) database (https://img.jgi.doe.gov/). Sequence alignment of 16S rRNA genes and LPAAT genes (also retrieved from IMG) were performed using Muscle and phylogenetic analyses were performed with MEGA7.0 (Kumar et al., 2016) with 500 bootstrap replicates. Sequence alignment was visualized using JalView (Waterhouse et al., 2009).

To search for SalA homologues in the *Tara* metagenome/metatranscriptomics datasets, we used the Ocean Gene Atlas (OGA) database OM_RGCv2_metaG (metagenomics) and OM-RGCv2_metaT (metatranscriptomics) with e-value cut-off of e^-40^ (Villar et al., 2018). Abundance was normalised as a percentage of the median mapped read abundance of genes/transcripts of ten prokaryotic single-copy marker genes (Milanese et al 2019). Taxonomic distribution of homologs was displayed using Krona in the OGA interface.

The genomes of the marine roseobacters used in this study were downloaded from the NCBI database. These comprised 9 strains that were found to produce SAL and 2 strains *(Stappia stellulata* DSM 5886 and *Dinoroseobacter shibae* DFL12) that did not. In order to identify genes potentially involved in SAL synthesis, each gene from the 11 genomes was assigned to an orthologous group using the eggNOG mapper (Huerta-Cepas, Forslund, *et al.*, 2016a). This program conducts a BLAST search of each sequence against the eggNOG database (Huerta-Cepas, Szklarczyk, *et al.*, 2016b) of orthologous genes, with the query sequence being annotated with the same orthologous group as the best BLAST hit. Orthologous groups that were present in the genomes of all SAL-producing strains but absent from the genomes of *S. stellulata* and *D. shibae* were considered to be potentially involved in SAL synthesis.

Abundance data of SalA homologues from four depths derived from the *Tara* metagenomics/metatranscriptomics datasets were tested for normal distribution using a Shapiro-Wilks test. Significant differences between depths was tested for using a Kruskal-Wallis test followed by a post-hoc Dunn’s test using Holm’s correction for multiple comparisons. All statistical analysis was performed in RStudio (version 1.3) using R (version 4.02).

### *In silico* homology modelling and docking studies for SalA

A SalA homology model was generated using the Phyre2 protein folding prediction server (Kelley et al. 2015) and the lyso-SAL lipid was drawn in MarvinSketch (v19.10.0, 2019, ChemAxon for Mac) and exported as a Mol SDF format file. The homology model was built using the structure of the lysophosphatidic acid acyltransferase PlsC (PDB code 5KYM, Robertson et al., 2017). The SalA protein model was then imported into Flare (v3.0, Cresset) for docking the lyso-SAL substrate and energy minimized with 2000 iterations with a cut off of 0.200 kcal/mol/A. The lyso lipid was imported as a ligand and energy minimized in Flare before being docked into the active site and the best scoring pose selected.

## Results

### A new sulfur-containing aminolipid is found in *Ruegeria pomeroyi* DSS-3

During LC-MS analysis of lipid extracts from *Ruegeria pomeroyi* DSS-3 grown on ½ YTSS medium, two prominent peaks eluting around 3.5 minutes were found in both negative and positive ionisation mode (**Figure 1**). The most prominent ions in the two peaks had *m/z* values of 656.6 and 672.7 in the negative ionisation mode, respectively. Other major lipids identified in this bacterium include two phospholipids, PG and PE and two aminolipids, ornithine lipid (OL) and glutamine lipid (QL) (Smith et al., 2019).

**Figure 1.**
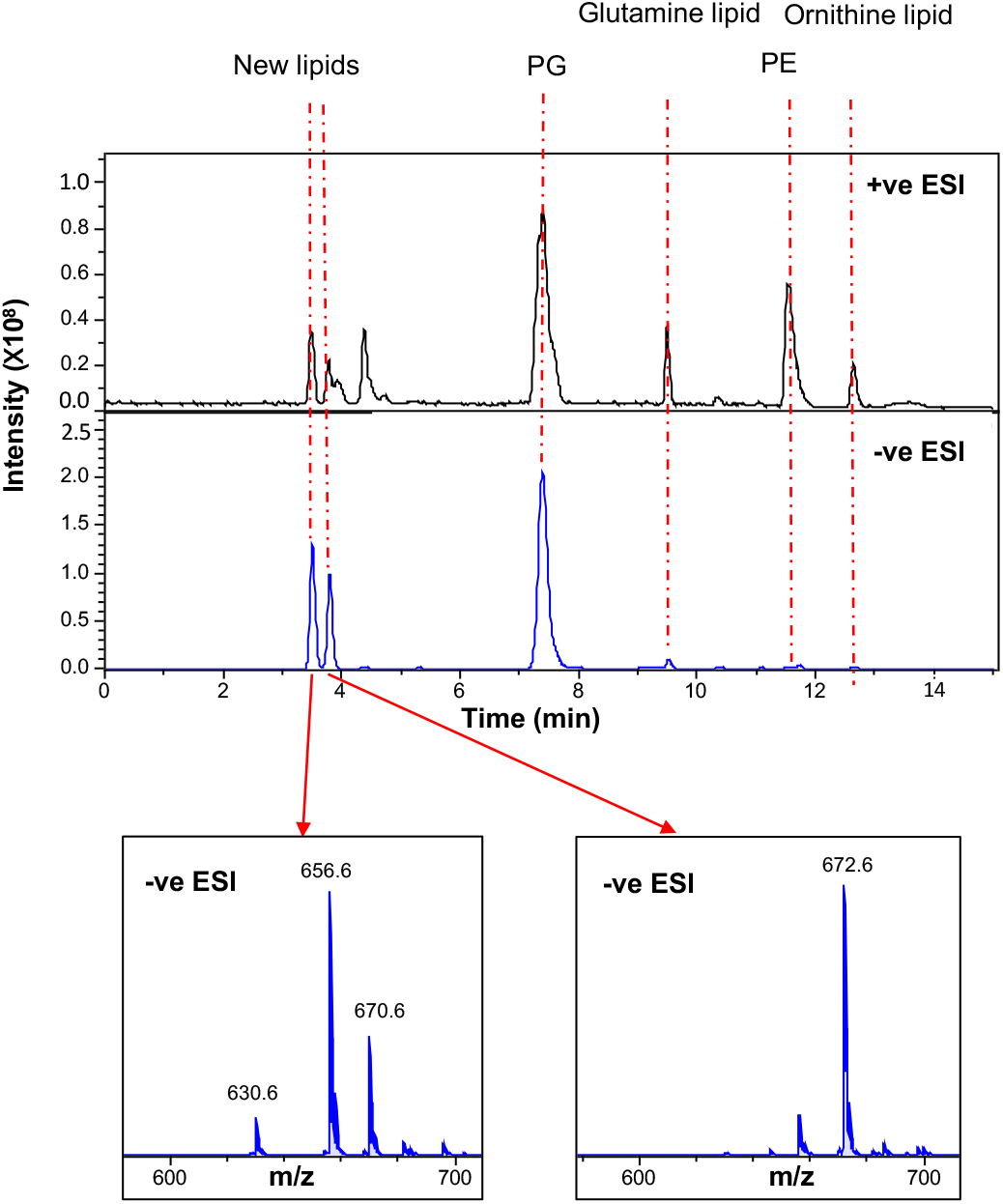
Lipid profiles of *Ruegeria pomeroyi* DSS-3 cultivated in 1/2 YTSS medium. MS spectra were obtained in both positive (+ve) and negative (-ve) ionisation mode using electrospray ionisation (ESI) using an amaZon SL ion trap MS (Bruker). PG, phosphatidylglycerol; PE, phosphatidylethanolamine. New lipids with *m/z* of 656.6 and 672.6 in -ve ESI were eluted between 3-4 min from the liquid chromatography column.

To elucidate the structure of the new lipids eluted at 3.5 min, the most intense species, at 656.4882 *m/z*, was selected for high resolution MS/MS analysis on a quadrupole - time of flight (Q-TOF) mass spectrometer (**Figure 2**). At low collision energy (40 eV) the major species formed corresponded to a neutral loss of 282 mass units. This is consistent with the neutral loss of an 18:1 fatty acid. A second peak at *m/z* 281.2480 is likely the carboxylate anion of an 18:1 fatty acid. Further fragmentation, at higher collision energies (up to 90 eV), yielded a major ion at *m/z* 237.2159. This ion likely corresponds to a 16:0 fatty acid present as a ketene, which would be consistent with the fragmentation scheme proposed for ornithine lipids and glutamine lipids (Zhang et al., 2009). These results therefore suggest a lipid class with a similar fatty acyl backbone structure to the aminolipids, such as ornithine and glutamine lipid (Smith et al., 2019). The glutamine lipid (QL, [M+H]^+^ *m/z* 719.7) and ornithine lipid (OL, [M+H]^+^ *m/z* 705.7) was eluted at 9.5 and 12.5 min, respectively (**Figure 1**). The formation of these novel lipids at ~3.5 - 4 min is not affected in the *olsA* or *glsB* mutants of *R. pomeroyi* DSS-3 (**Supplementary Figure S1**). The *olsA* and *glsB* genes in *R. pomeroyi* DSS-3 were essential for the production of the nitrogen-containing ornithine/glutamine lipids (Smith et al., 2019)

**Figure 2.**
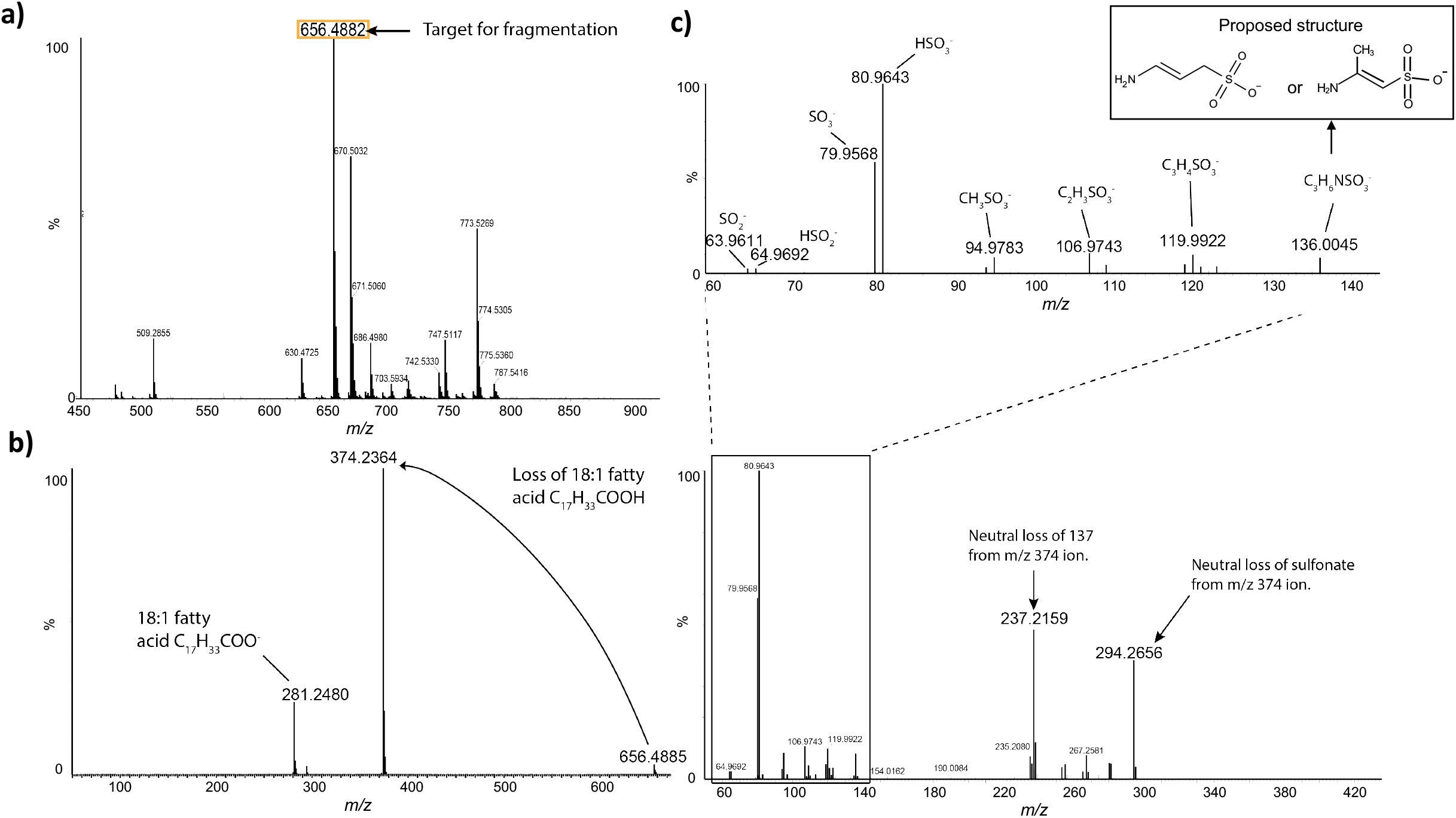
A previously unidentified class of sulfur-containing aminolipids (SAL) is present in *Ruegeria pomeroyi* DSS-3. **a)** Intact masses of *R. pomeroyi* lipids, measured using a high resolution, accurate mass quadrupole-time of flight (Q-TOF) mass spectrometer (Waters Synapt G2-Si) in negative ionisation mode. The identity of the most abundant ion, highlighted, was unknown, so this ion was fragmented in order to elucidate its structure. **b)** Fragmentation spectrum of *m/z* 656.4882 ion at 40 eV collision energy (MS^2^). The most abundant species corresponds to the loss of a 18:1 fatty acid. **c)** Fragmentation spectrum of *m/z* 3742364 ion at 90 eV collision energy (MS^3^). The lower spectrum shows a peak at 237.2159, consistent with the presence of a 16:0 fatty acid. The upper spectrum shows an expanded view of the spectrum in the *m/z* range below 140. Ions are annotated with their predicted elemental composition. Inset is the proposed structure of the 136.0045 *m/z* fragment (3-aminopropane sulfonic acid or 2-aminopropane sulfonic acid, respectively). *Ruegeria pomeroyi* DSS-3 was cultivated in 1/2 YTSS medium.

Prominent peaks at 80 and 81 *m/z*, respectively, were apparent in the fragmentation spectrum obtained at 90 eV collision energy (**Figure 2c**). The accurate masses of these ions were 79.9568 and 80.9643. Of the candidate formulae within 100 ppm of the measured mass, 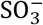 and 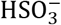 appear most plausible, with mass errors of 0.182 ppm and 4.194 ppm, respectively (**Table 1**). A smaller peak doublet at *m/z* 63.9611 and 64.9692 was also present in the 90 eV spectrum. These masses are unambiguously assigned to 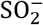 (mass error 12.506 ppm) and 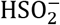 (mass error 8.08 ppm). Taken together, these results demonstrate the presence of a sulfonate group in the lipid. An ion at 136.0045 *m/z* corresponded to the deprotonated head group. The mass determined here is larger than that of deprotonated taurine (*m/z* 124). Since the head group includes a sulfonate (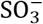) group, the plausible formula most closely corresponding to the accurate mass is C_3_H_6_NSO_3_ (**Table 1**). This is consistent with the structure being aminopropane sulfonic acid although the position of the amino group cannot be unequivocally determined by mass spectrometry (**Figure 2**). The proposed fragmentation scheme is presented in **Figure 3**.

**Figure 3.**
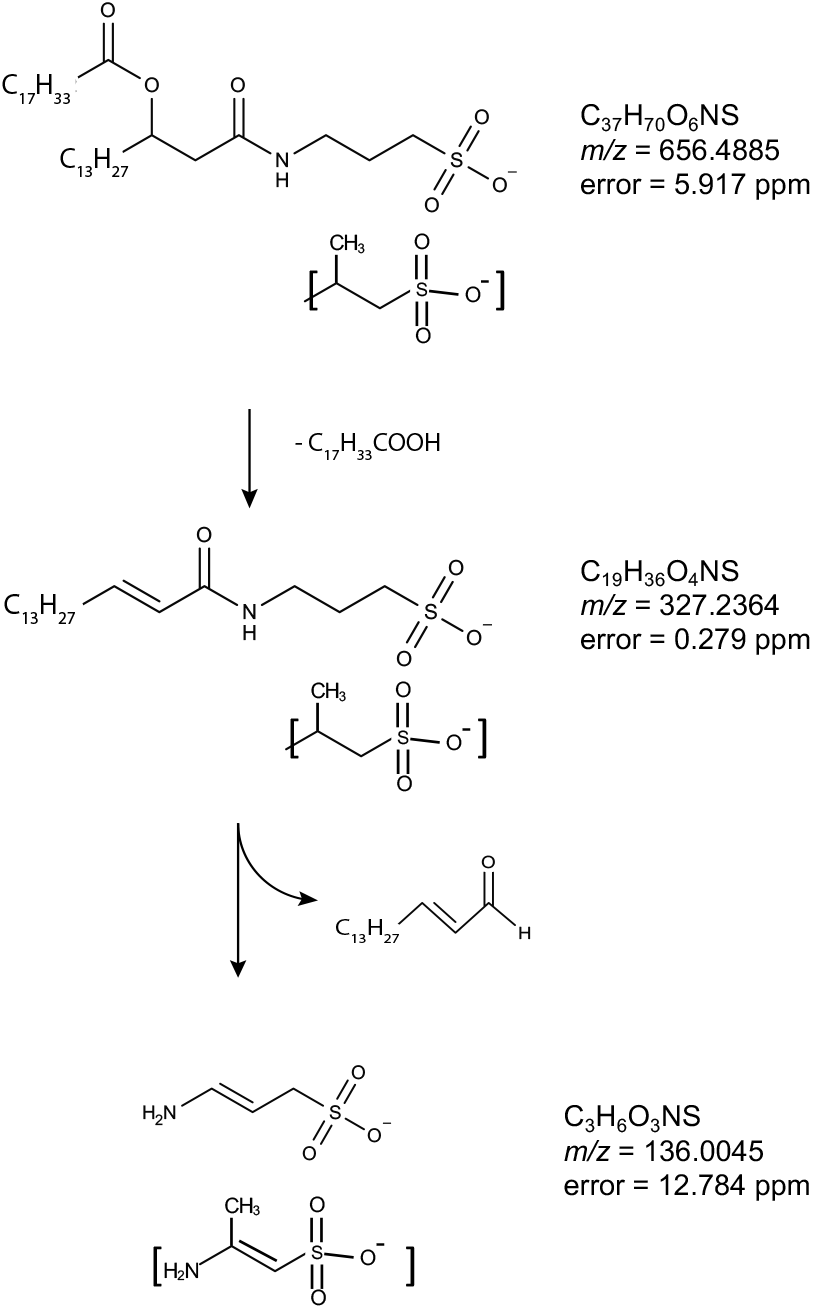
Proposed fragmentation scheme for the SAL lipid with *m/z* 656.4885. Mass errors indicate the difference between the measured and theoretical masses of the proposed species. The aminolipid head group is consisted of an aminopropane sulfonic acid of 3-aminopropane sulfonic acid (a.k.a. homotaurine) or 2-aminopropane sulfonic acid (as indicated by square brackets).

**Table 1.**
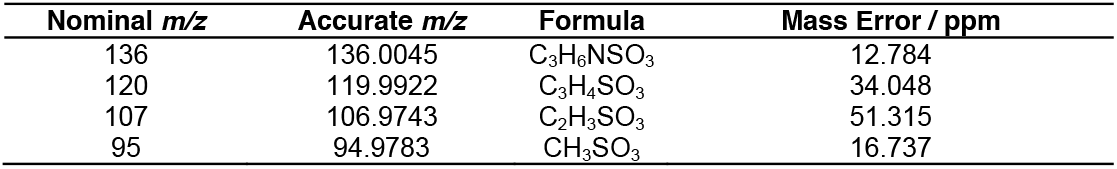
Nominal and accurate masses of proposed head group fragments, with mass errors implied by the proposed formulae.

| Nominal $m/z$ | Accurate $m/z$ | Formula | Mass Error / ppm |
| --- | --- | --- | --- |
| 136 | 136.0045 | C<sub>3</sub>H<sub>6</sub>NSO<sub>3</sub> | 12.784 |
| 120 | 119.9922 | C<sub>3</sub>H<sub>4</sub>SO<sub>3</sub> | 34.048 |
| 107 | 106.9743 | C<sub>2</sub>H<sub>3</sub>SO<sub>3</sub> | 51.315 |
| 95 | 94.9783 | CH<sub>3</sub>SO<sub>3</sub> | 16.737 |

To further confirm the presence of an amino-group in the hydrophilic head of this SAL, we cultivated *Ruegeria pomeroyi* DSS-3 in a chemically defined marine ammonium mineral salts (MAMS) medium using ^15^N-ammonium as the sole nitrogen source. Indeed, the ^15^N-labelled SAL was readily observed in the lipid extract resulting in a shift of *m/z* from 656.4951 to 657.4903 (**Supplementary Figure S2a**) whereas the non-nitrogen containing lipids, such as PG were not labelled by ^15^N as expected (**Supplementary Figure S2b**). The incorporation of the ^15^N isotope into the head group of SAL was confirmed by MS^n^ (**Supplementary Figure S2c, d**). We also performed the same MS^n^ analysis on the *m/z* 672.4875 species as well as the ^15^N-labelled *m/z* 673.4852 species. Loss of 282 at MS^2^ (672.4875--> 390.2317; 673.4852-->391.2285) suggests the R_2_ fatty acid was C18:1. Therefore, the data suggest that the lipid species eluted immediately after the *m/z* 656.6 species is likely a hydroxylated SAL and the proposed fragmentation scheme is presented in **Supplementary Figure S3**.

### The sulfur-containing aminolipid is found in a range of marine roseobacters

To investigate the presence of SAL amongst roseobacters we selected 16 strains, in addition to *R. pomeroyi* DSS-3, to obtain a wide coverage of the roseobacter group including the model roseobacter bacterium *Phaeobacter inhibens* DSM 17395 (**Figure 4**). The selected strains included *Stappia stellulata*, which recent phylogenetic studies indicate is not a member of the *Rhodobacteraceae* (Pujalte et al., 2013), which served as an outgroup. These strains were each grown in marine broth overnight, before cells were harvested for lipid analysis. SAL was detected in all the strains tested apart from *S. stellulata* and *Dinoroseobacter shibae* (**Figure 4a**). The separation of these two strains from the remaining roseobacter sequences is in line with previous results showing *D. shibae* branching deeply within the *Rhodobacteraceae* phylogeny (Simon et al., 2017).

**Figure 4.**
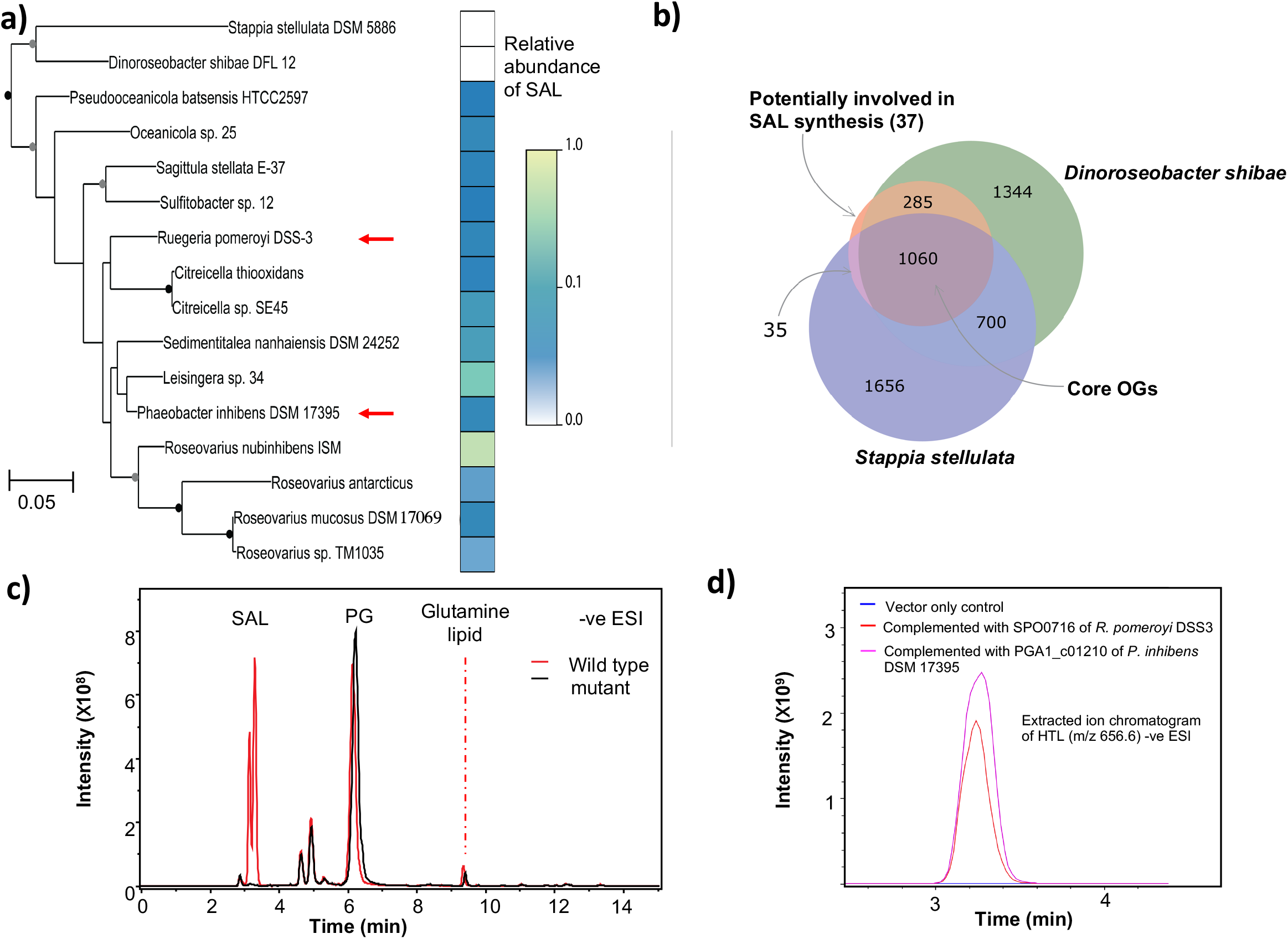
**a)** Phylogeny of 16S rRNA gene sequences from selected *Rhodobacteraceae* plotted alongside information on the abundance of the SAL. Nodes with more than 50% bootstrap support are indicated with circles (grey: 50 - 69 %; black: > 70%). The red arrows point to the two bacteria which were used as models to investigate SAL biosynthesis. The *Stappia stellulata* 16S rRNA gene sequence was used as an outgroup. The abundance of SAL measured in each strain, as a proportion of the maximum abundance measured, is plotted alongside the phylogeny. One ml culture of OD_540_~1.0 was collected by centrifugation before lipid extraction and abundances were calculated as the peak area of SAL species relative to a phosphatidylglycerol internal standard (0.25 μM diheptadecanoyl phosphatidylglycerol, C17/C17 PG). **b)** Identification of two candidate acyltransferases to be investigated for involvement in SAL biosynthesis. The Venn diagram illustrates the logic used to identify the candidate genes. The central circle represents the 1417 orthologous groups (OGs) common to all SAL-producing strains. **c)** LC-MS chromatogram showing the lipid profile of wild type and the *salA* mutant of *Phaeobacter inhibens* DSM 17395. The absence of SAL lipids in the mutant is evident in the black trace. **d)** Extracted ion chromatogram (EIC) showing the complementation of the *salA* mutant of *Phaeobacter inhibens* DSM 17395 by either *salA* of *R. pomeroyi* DSS3 (SPO716) or *P. inhibens* DSM 17395 (PGA1_c01210) which restores the production of SAL *(m/z* 656.6). The empty vector control is shown in blue. Bacterial cells were cultivated in marine broth medium. Ions were analysed in negative (-ve) ionisation mode using electrospray ionisation (ESI) using an amaZon SL ion trap MS (Bruker)

### Comparative genomics to determine genes involved in SAL biosynthesis

We then conducted a comparative genomics investigation into the roseobacter strains whose lipid profiles had been analysed. We reasoned that synthesis of the SAL would require an *N*-acyltransferase activity to acylate aminopropane sulfonic acid, analogous to that mediated by OlsB and GlsB in the synthesis of ornithine and glutamine lipid (Gao et al., 2004; Smith et al., 2019). We investigated predicted *N*-acyltransferases that were present in all the strains that produced SAL in marine broth (the ‘producers’) while being absent from the strains that did not produce SAL (the ‘non-producers’). We assigned all the genomic sequences from the 9 genome-sequenced producer strains and 2 non-producer strains to orthologous groups (OGs) using the eggNOG-mapper software (Huerta-Cepas, Forslund, et al., 2016a), which provides a consistent pipeline for sequence annotation and OG assignment by comparison to the eggNOG database (Huerta-Cepas, Szklarczyk, et al., 2016a). We identified a group of 1417 ‘core’ genes which were present in the genomes of all SAL producer strains of which 1060 were also present in the two non-producers (**Figure 4b**). 37 candidate genes are present in all SAL producer strains but not in the genomes of the non-producers (**Figure 4b**), two of which (OG accession numbers 08UX5 and 05CDD) were annotated as being potential acyltransferases (**Table 2**). We therefore generated mutants in these two genes in the two model bacteria, *R. pomeroyi* DSS-3 and *P. inhibens* DSM 17395 and screened for the loss of SAL production. The 08UX5 mutant (locus SPO2471) of *R. pomeroyi* DSS-3 still produced SAL to the same level as the wild type (data not shown), suggesting that this gene is unlikely involved in SAL formation. However, in the 05CDD mutant of *P. inhibens* DSM 17395 (locus tag PGA1_c01210), SAL formation is completely abolished, suggesting that this gene is indeed responsible for SAL biosynthesis (**Figure 4c**). This gene is named *salA* hereafter. Indeed, when the mutant was complemented with either *salA* from *R. pomeroyi* DSS-3 (SPO0716) or *P. inhibens* DSM 17395 (PGA1_c01210), SAL production was restored (**Figure 4d**).

**Table 2.** Candidate eggNOG orthologous groups (OGs) that are unique in the genomes of SAL-producers. The two predicted acyltransferases are highlighted in bold.

| OG | Locus tag | COG category | eggNOG annotation |
| --- | --- | --- | --- |
| 07TNN | SPO0365 | S | Domain of unknown function (DUF1989) |
| **05CDD** | **SPO0716** | **S** | **Phospholipid glycerol acyltransferase** |
| 01VSN | SPO0838 | M | Bacterial sugar transferase |
| 05MEE | SPO0968 | L | Nudix Hydrolase |
| 05K39 | SPO1109 | S | Endonuclease Exonuclease phosphatase |
| 05NU0 | SPO1287 | S | Glyoxalase bleomycin resistance protein dioxygenase |
| 08EFD | SPO1424 | P | Sodium hydrogen exchanger |
| 08K1F | SPO1437 | I, Q | Dehydrogenase |
| 05D82 | SPO1743 | E | Glutamate dehydrogenase |
| 05S99 | SPO1919 | P | Tellurite resistance protein |
| 090P8 | SPO1963 | G | Phosphoglycerate mutase |
| 05N59 | SPO2284 | M | Transglycosylase |
| 01QFF | SPO2332 | P | Acriflavin resistance protein |
| 08YCB | SPO2344 | E | Sarcosine oxidase, gamma subunit |
| **08UX5** | **SPO2471** | **S** | **Acetyltransferase, (GNAT) family** |
| 01QH6 | SPO2697 | C | CoA-binding domain protein |
| 07QRG | SPO2741 | L | DNA polymerase iii |
| 05X3J | SPO2754 | S | Uncharacterized protein |
| 060TV | SPO2757 | S | EF hand domain protein |
| 01TDR | SPO2815 | P | ABC transporter permease protein |
| 08TCB | SPO3054 | S | Uncharacterized protein |
| 0808N | SPO3431 | S | Domain of unknown function (DUF3576) |
| 08RQB | SPO3439 | I | Enoyl-coA hydratase isomerase family protein |
| 08R76 | SPO3444 | M | 3-deoxy-d-manno-octulosonic-acid transferase |
| 07RFB | SPO3740 | O | Catalyses the reversible oxidation-reduction of methionine sulfoxide in proteins to methionine |
| 6405 | SPO0725 | S | SH3 type 3 |
| 01RKE | SPO2816 | P | ABC transporter permease protein |
| 05DM9 | SPO0035 | S | Core-2 I-branching enzyme family protein |
| 0627S | SPO_RS14095 | S | Uncharacterized protein |
| 07SF7 | SPO2814 | E | Peptide opine nickel uptake family ABC transporter periplasmic substrate-binding protein |
| 05SH6 | SPO3502 | S | DUF3118 protein |
| 08U9J | SPO3819 | S | Type I secretion target repeat protein |
| 05WTG | SPO2776 | S | Uncharacterized protein |
| 05Q6E | SPO3716 | S | Uncharacterized protein |
| 08T58 | SPO3836 | S | Mg<sup>2+</sup> transport protein CorA |
| 064YK | SPO1241 | S | Uncharacterized protein |
| 05U51 | SPO0776 | S | Thioesterase |

SalA is a putative O-acetyltransferase-like protein with a recognized LPAAT (lysophosphatidic acid acyltransferase) domain. Amongst bacterial LPAAT-domain containing proteins, the best characterized examples are PlsC and OlsA, encoding enzymes responsible for the final step in the biosynthesis of the anionic phospholipid phosphatidic acid (PA) and the ornithine/glutamine-containing aminolipid, respectively (Smith et al., 2019; Röttig & Steinbüchel, 2013, Korbes et al., 2016). The structure of PlsC has recently been solved, showing an *in silico* docked LPA lipid together with the fatty acid in an acyl carrier protein (ACP, Robertson et al., 2017). Multiple sequence alignments of SalA, PlsC and OlsA shows the presence of two conserved sequence motifs (**Figure 5**), representing the catalytic centre (HX_4/5_D) and the substrate co-ordination centre (FP[E/S]G[T/V]), respectively. Notably, both PlsC and OlsA have the conserved HX_4_D motif whereas SalA has the HX_5_D motif. Interestingly, the reported key Lys105 in PlsC, thought to be responsible for electrostatic interactions via its amide nitrogen backbone to the negatively-charged oxygen of the ACP-fatty acid intermediate, was replaced with Arg135 in SalA. The LPA phosphate head group is thought to be coordinated by Arg159 in PlsC. However, the sequence alignment shows a Val189 in SalA. In order to further investigate the implications of the sequence alignment, we obtained a homology model of SalA. The model shows the catalytic HX_5_D motif to be structurally comparable to that of PlsC despite the additional residue, with the His109 and Asp115 adjacent to each other, analogous to that in PlsC (**Supplementary Figure S4**). *In silico* docking of a lyso-SAL lipid molecule into the model demonstrated a possible pose for the lyso-lipid hydroxy group adjacent to His109 (**Supplementary Figure S4**) with the Arg135 suggested to coordinate the sulfonate head group. The conformationally flexible alkyl chain group was able to adopt many configurations but the polar head group was docked consistently in the same region. Overall, the data suggests a diversification in function of LPAAT family enzymes during evolution, with SalA representing a novel member of this group. The presence of this unique motif of HX_5_D in SalA allowed us to determine the distribution of SAL-biosynthesis in environmental metagenomes and metatranscriptomes (see below).

**Figure 5.**
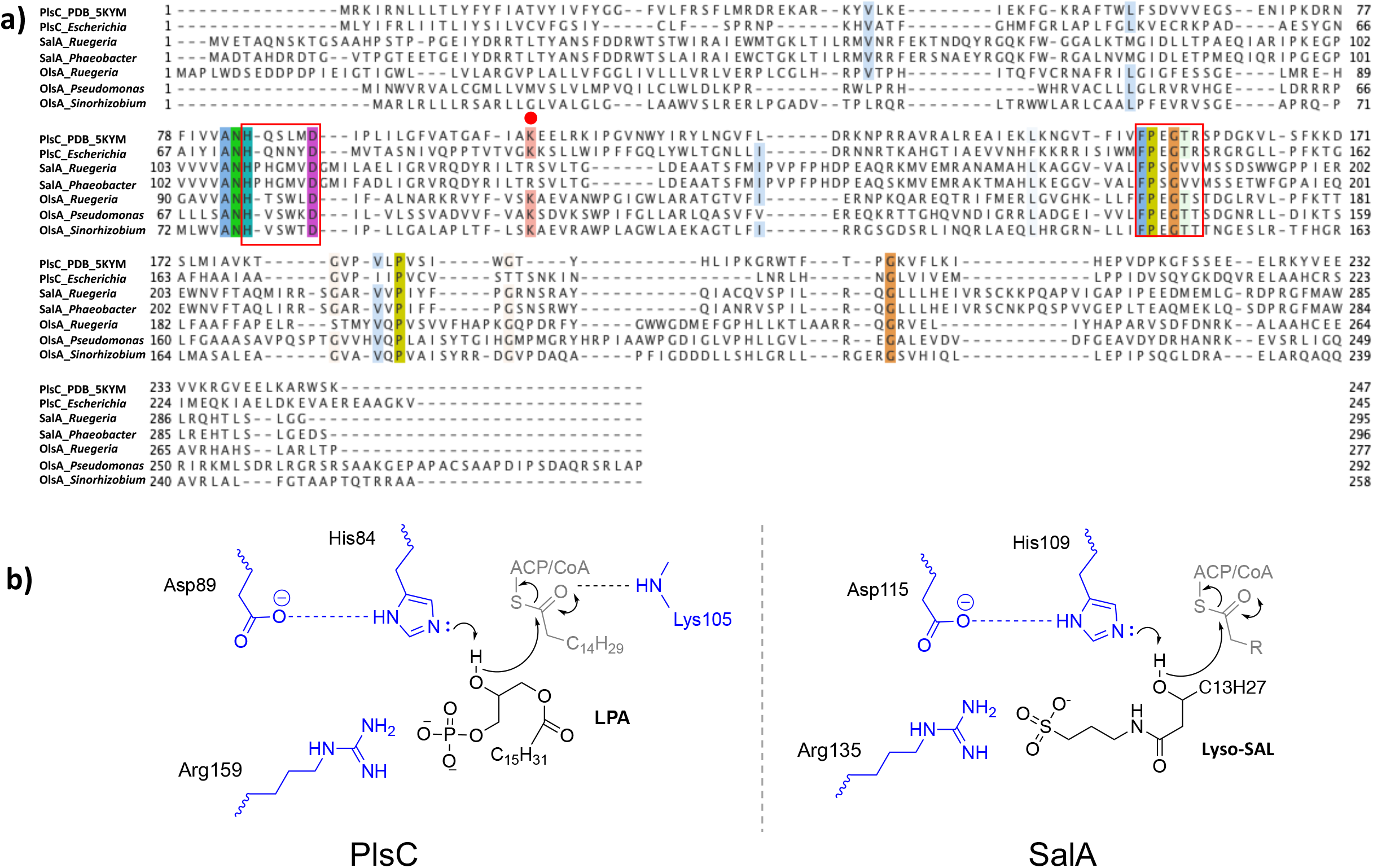
SalA represents a new member of the lysophosphatidic acid acyltransferase (LPTAA) family. **a)** Multiple sequence alignment of SalA from *Ruegeria pomeroyi* DSS-3 and *Phaeobacter inhibens* DSM 17395, lysophosphatidic acid acyltransferase PlsC from *Escherichia coli* and *Thermotoga maritima* (PDB code 5KYM, Robertson et al., 2017), and OlsA involved in ornithine lipid biosynthesis from *R. pomeroyi* DSS-3 (SPO1979, Smith et al., 2019), *Pseudomonas aeruginosa* (PO4351, Lewenza et al., 2011) and *Sinorhizobium meliloti* (Weissenmayer et al., 2002). The two red boxes highlight the catalytic centre (HX_n_D) and the substrate binding site (FPXGXX), respectively. The red dot represents the key lysine 105 involved in catalysis in PlsC. **b)** Proposed residues involved in PlsC and SalA catalysis. LPA, lysophosphatidic acid.

### SAL production in *Phaeobacter inhibens* DSM 17395 is involved in biofilm formation

We next investigated the role of SAL lipids in the physiology of roseobacters. The loss of SAL lipids had no clear role in the growth of the bacterium. Both wild type and the *salA* mutant of *Phaeobacter inhibens* DSM 17395 had comparable growth rates and reached similar final cell density in marine broth medium (**Supplementary Figure S5a**). An important change of lifestyle for roseobacters is the switch from planktonic growth to biofilm formation which triggers a particle-associated life strategy that is ecologically relevant for their survival in the natural environment (Sule & Belas 2013). It has been shown previously that many roseobacters including *Phaeobacter inhibens* DSM 17935 are able to form a biofilm and a 65 kb plasmid in this bacterium was important for biofilm formation (Frank et al., 2015; Michael et al., 2016). Interestingly, we observed that the *salA* mutant has a significantly reduced ability to form biofilms when in contact with solid surfaces such as glass (**Figure 6**) and plastics (**Supplementary Figure S5b**). Both the biofilm on the glass surface as well as the thickness of the biofilm are significantly reduced in the *salA* mutant strain in the early phase of biofilm formation (3 h) and the latter stage (24 h, 48 h) of biofilm maturation (**Figure 6**). The 65kb biofilm plasmid was confirmed to be present in the *salA* mutant (**Supplementary Figure S5c**). Thus, the significant reduced ability of the *salA* mutant in biofilm formation suggests that this lipid may play a key role in roseobacters in their natural environment.

**Figure 6.**
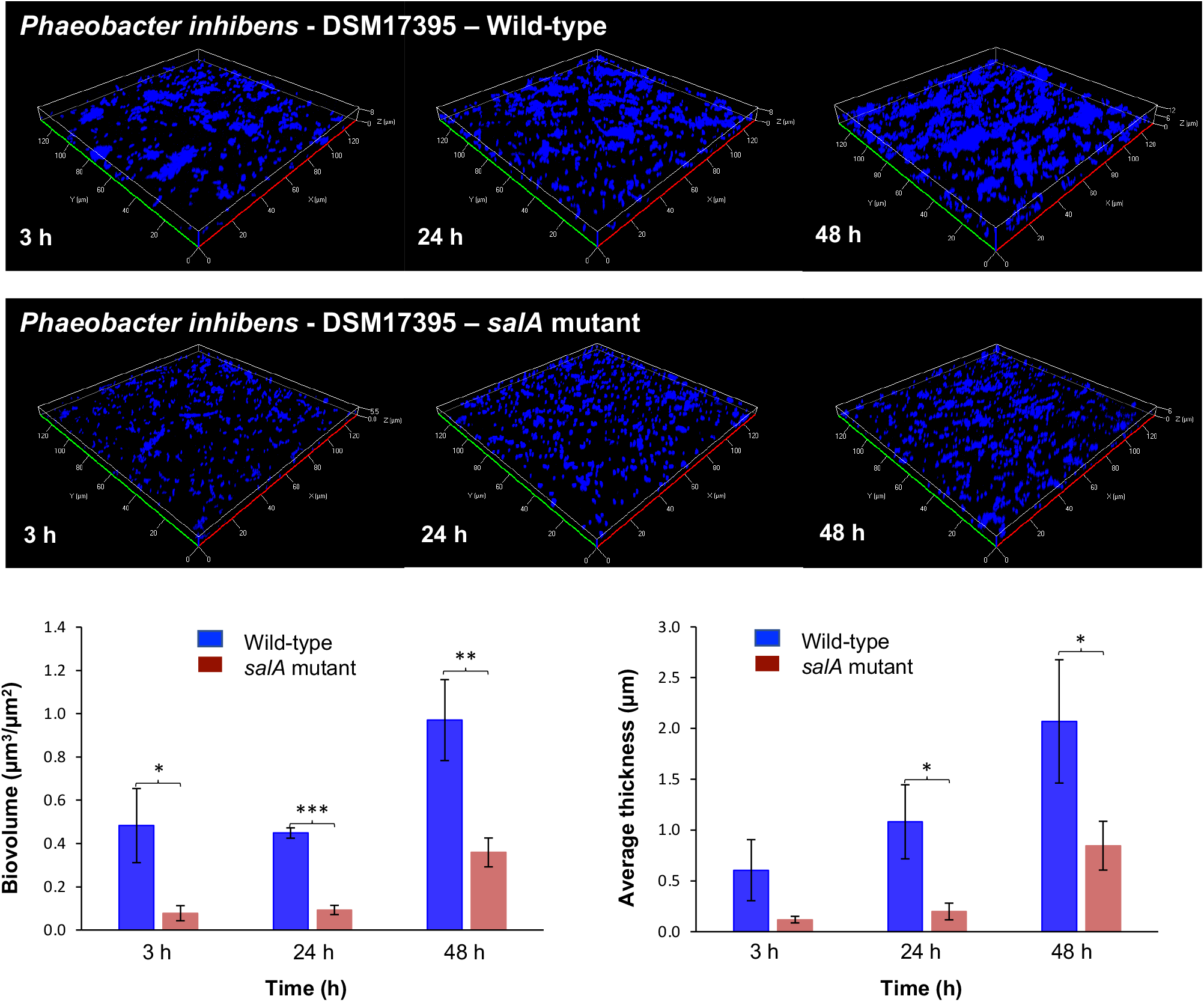
Quantification of biofilm formation in *Phaeobacter inhibens* DSM 17395 wild type and the *salA* mutant. Biofilm was developed for 3 h, 24 h and 48 h respectively. The biovolume and the thickness of the biofilm over time were quantified. *, *p*<0.05, **, *p*<0.01, ***, *p*<0.001. Bacterial cells were cultivated in marine broth medium.

### Distribution of the new acetyltransferase SalA in the *Tara* Ocean metagenomes and metatranscriptomes

To better understand the distribution of SAL in environmental microbial assemblages, we searched the *Tara* Ocean metagenomes and metatranscriptomics datasets using SalA (locus tag, SPO0671 of *R. pomeroyi* DSS-3) as the query. We experimentally determined the e value cut-off to be e^-40^ at which value it selectively retrieves LPAAT homologues belonging to SalA but not OlsA or PlsC. The environmental SalA homologues obtained from the *Tara* Oceans metagenome and metatranscriptomics dataset were aligned and the key sequence motifs were manually examined. In particular, the HX_5_D motif is strictly conserved in all SalA sequences retrieved from the *Tara* Oceans datasets, providing strong support that these environmental sequences are of the SalA but not PlsC nor OlsA clade. On average, between 2-4% of microbial cells are estimated to have the potential for SAL biosynthesis; this is comparable to that of the *olsA* gene but somewhat lower than the PlcP gene in the same dataset, suggesting SAL biosynthesis is less prevalant than the PlcP-mediated lipid remodelling pathway (Sebastian et al., 2016, Smith et al., 2019). This is likely due to the fact that SALs are primarily found in marine roseobacters but not in other dominant marine *Alphaproteobacteria*, such as the abundant bacterium *Pelagibacter ubique* of the SAR11 clade which are capable of PlcP-mediated lipid remodelling (Carini et al., 2015; Sebastian et al., 2016). Indeed, the majority (>85%) of the SalA sequences from the *Tara* Oceans dataset were classified as members of the *Rhodobacteraceae* in both *Tara* Oceans metagenomes and metatranscriptomes (**Figure 7**), and a thorough search of 120-genome sequenced *Rhodobacteraceae* confirmed the wide occurrence of *salA* in all 10 clades of the roseobacters (**Supplementary Figure S6**, Bartling et al., 2018).

**Figure 7.**
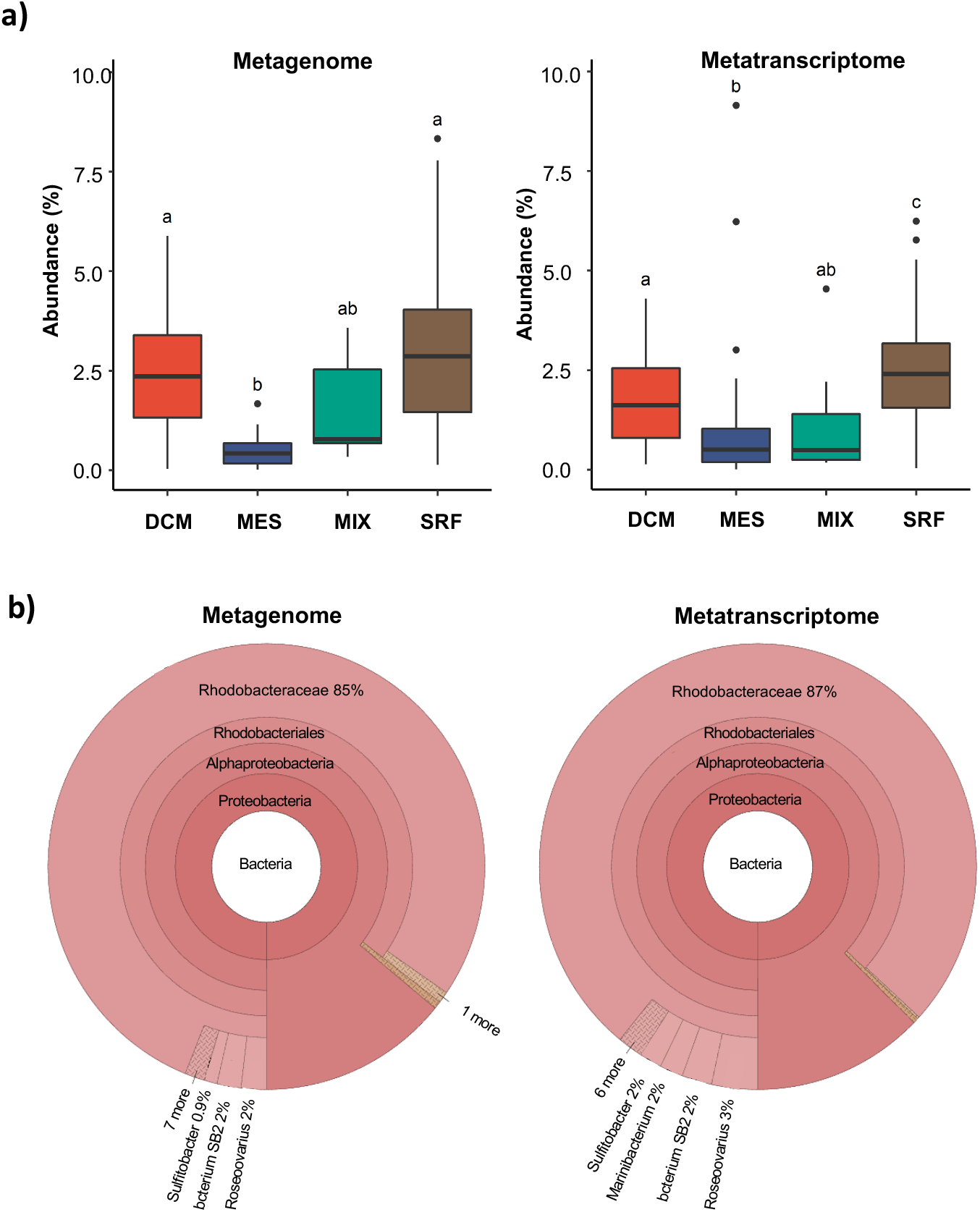
The distribution of the *salA* gene in the *Tara* oceans database. **a)** The Ocean Gene Atlas (OGA) database was searched using *salA* of *Ruegeria pomeroyi* DSS-3 (SPO0716) with an e-value cut-off of e^-40^ and the abundance was calculated as a percentage of median abundance of ten prokaryotic single-copy marker genes/transcripts (Villar et al., 2018). **b)** The taxonomic distribution of homologs was displayed using Krona in the OGA interphase. DCM, deep chlorophyll maximum; SRF, surface water; MES, mesopelagic zone; MIX, mixed layer. More than 85% of the SalA homologs are classified as marine roseobacters (*Rhodobacteraceae*) in metagenome (left panel) and metatranscriptome (right panel).

## Discussion

Here, we identify a novel aminolipid containing an aminopropane sulfonic acid head group that is widespread amongst marine roseobacters. The presence of a sulfonate group means this SAL lipid also falls into the broad category of sulfonolipids. The most abundant, and arguably one of the best studied lipids of this type, is sulfoquinovosyl diacylglycerol (SQDG) which is present in the membranes of most oxygenic phototrophs (Frentzen, 2004) as well as some heterotrophic bacteria (Villanueva et al., 2013). SQDG likely plays a structural role in photosynthetic membranes, since crystal structures of photosystem proteins show specific binding of this lipid (Umena et al., 2011).

Other sulfolipids appear to elicit potent responses when certain organisms are exposed to them. Thus, a sulfolipid produced by zooplankton from a number of copepod species was found to induce toxin production in the dinoflagellate *Alexandrium minutum* (Selander et al., 2015), likely as a defence against predation. Conversely, a sulfonolipid produced by the *Bacteroidetes* bacterium *Algoriphagus machipongonensis* induced the development of multicellularity in a choanoflagellate (Alegado et al., 2012). Both examples suggest that sulfolipids are used by the sensing organism as a marker for the presence of another organism with which it interacts (either as a predator or as a symbiont). The fact that sulfolipids appear to be relatively rare across the tree of life likely makes them well suited to mediate such chemical interactions where a high degree of specificity is required. Lipids similar to those produced by *A. machipongonensis* have been described in a number of *Bacteroidetes*, particularly amongst *Cytophaga* (Godchaux & Leadbetter, 1980, 1983, 1984). They tend to be localised to the outer membrane, and seem to play a role in the gliding motility of these organisms (Abbanat et al., 1986; Godchaux & Leadbetter, 1988). The sulfonolipids from *Bacteroidetes* differ from those that we describe here in roseobacters in that they are composed of a base, termed capnine, similar to the sphingoid bases of sphingolipids, which may be *N*-acylated to form the full sulfonolipid (Godchaux & Leadbetter, 1980). In this way they are similar structurally to sphingolipids whereas the SALs of the roseobacter group are more similar to aminolipids such as ornithine lipid and glutamine lipid (**Figure 5**). Whether the SAL lipid plays a role in interspecies interactions requires further work. However, we already observed that this lipid is involved in biofilm formation in *Phaeobacter inhibens* DMS17395 (**Figure 6**), suggesting that formation of this SAL lipid may play an important role in the adaptation of marine roseobacters to a biofilm lifestyle.

A survey of the distribution of SAL among isolates from the roseobacters indicated that the ability to produce this lipid is widely distributed within the group. One strain, *D. shibae*, taxonomically the most basal of the strains examined, lacked any SAL under the conditions assessed, as did the outgroup strain *Stappia stellulata.* The absence of SAL in these strains suggests they lack the capacity to produce this lipid as the other roseobacters examined seem to produce SAL constitutively. However, it is possible that these strains have the capacity to produce SAL, but only do so under certain conditions. This pattern is observed for ornithine lipid, which is produced constitutively in some bacteria, such as *R. pomeroyi* DSS-3 (Smith et al., 2019), but in others is only produced as a response to P-depletion (Carini et al., 2015; Minnikin & Abdolrahimzadeh, 1974). Indeed, a close *salA* homolog was found in the genome of *D. shibae* (Dshi_0206), but it is absent in *S. stellulata*.

Although we have identified the LPAAT enzyme, SalA, involved in the last step of synthesis of this new sulfur-containing aminolipid, the key steps and genes involved in the synthesis of the lyso-SAL lipid remain to be determined. It is likely that SAL synthesis occurs in a manner analogous to that of ornithine and glutamine lipids. As such, 3-hydroxy fatty acids would be required as a substrate for the first step in SAL synthesis (Weissenmayer et al., 2002). Such a hypothesis suggests that the aminopropane sulfonic acid moiety is also directly produced by the marine roseobacters since no exogenous supply was provided. The presence of 3-aminopropane sulfonic acid (a.k.a. homotaurine) has been documented in some red algae (Ito et al., 1977; Miyasawa et al., 1970) and unicellular green algae (prasinophytes such as *Ostreococcus* and *Micromonas*, Durham et al., 2019) but, to the best or our knowledge, never previously in bacteria. However, a hydroxylated form of 2-aminopropane sulfonic acid, cysteinolic acid, has been found in a variety of marine phytoplankton and heterotrophic bacteria, including *Ruegeria pomeroyi* DSS-3 although its biosynthetic pathway remains to be established (Durham et al., 2019). Nevertheless, it is tempting to speculate that 2-aminopropane sulfonic acid is likely the hydrophilic head of the new SAL observed in these marine roseobacters and this certainly warrants further investigation.

As a structural analogue of γ-aminobutyric acid (GABA), aminopropane sulfonate such as homotaurine appears to activate GABA-receptors in the mammalian brain, a property which has led to its proposal as a potential therapeutic for Alzheimer’s disease (Aisen et al., 2011). Enzymes involved in the catabolism of GABA also show similar levels of activity against homotaurine as they do with GABA as a substrate (De Gracia & Jollès-Bergeret, 1973; Mayer & Cook, 2009). Indeed, several roseobacters, including *R. pomeroyi* DSS-3, are able to use homotaurine as a sole nitrogen source (Mayer & Cook, 2009). The relative availability of GABA and homotaurine in environments such as seawater has not been studied so it remains unclear which of these compounds is the primary substrate of the enzymes capable of their catabolism in the natural environment. Questions remain as to how roseobacters, such as *R. pomeroyi* DSS-3, synthesize aminopropane sulfonate to be used in the formation of the SAL lipid. It is known, however, that the biosynthesis of taurine, a structural analogue of homotaurine, occurs via oxidation of cysteine to cysteine sulfinic acid by a cysteine dioxygenase, followed by decarboxylation to yield hypotaurine and eventually taurine (Agnello et al., 2013). A homologue of cysteine dioxygenase is indeed found in *Ruegeria pomeroyi* DSS-3 (locus tag, SPO3687). However, a deletion mutant of SPO3687 still produced SAL (data not shown). Thus, its role in homotaurine biosynthesis is questionable. Therefore, the genes underpinning the biosynthesis of the lyso-SAL in roseobacters remain to be identified.

To sum up, this study describes a new class of lipid, which are an important component of the membranes of a number of marine *Rhodobacteraceae*. Comparative genomics of SAL-producing strains has identified a novel acyltransferase (SalA) which is involved in the production of this lipid. *salA* is widely distributed in marine microbial assemblages in the Oceans and actively expressed in *Tara* Oceans metatranscriptomes and its functional role in addition to biofilm formation in these marine bacteria certainly warrants further investigation.

## Acknowledgements

This project has received funding from the European Research Council (ERC) under the European Union’s Horizon 2020 research and innovation programme (grant agreement no. 726116) and the Deutsche Forschungsgemeinschaft (DFG, German Research Foundation) - Project-ID 34509606 - TRR 51. We also thank the Natural Environment Research Council, UK. for funding the PhD studentship awarded to AFS. We acknowledge Dr I. Hands-Portman and the Imaging Suite at the School of Life Sciences, University of Warwick for providing assistance with the use of confocal microscopy. We are grateful to Prof J.D. Todd, Dr H. Schäfer, Dr A. Curson, Prof M. Simon and Dr H.-A. Giebel for many insightful discussions.

## Classification

Microbial ecology and functional diversity of natural habitats

## Supplementary information

**Table S1.**
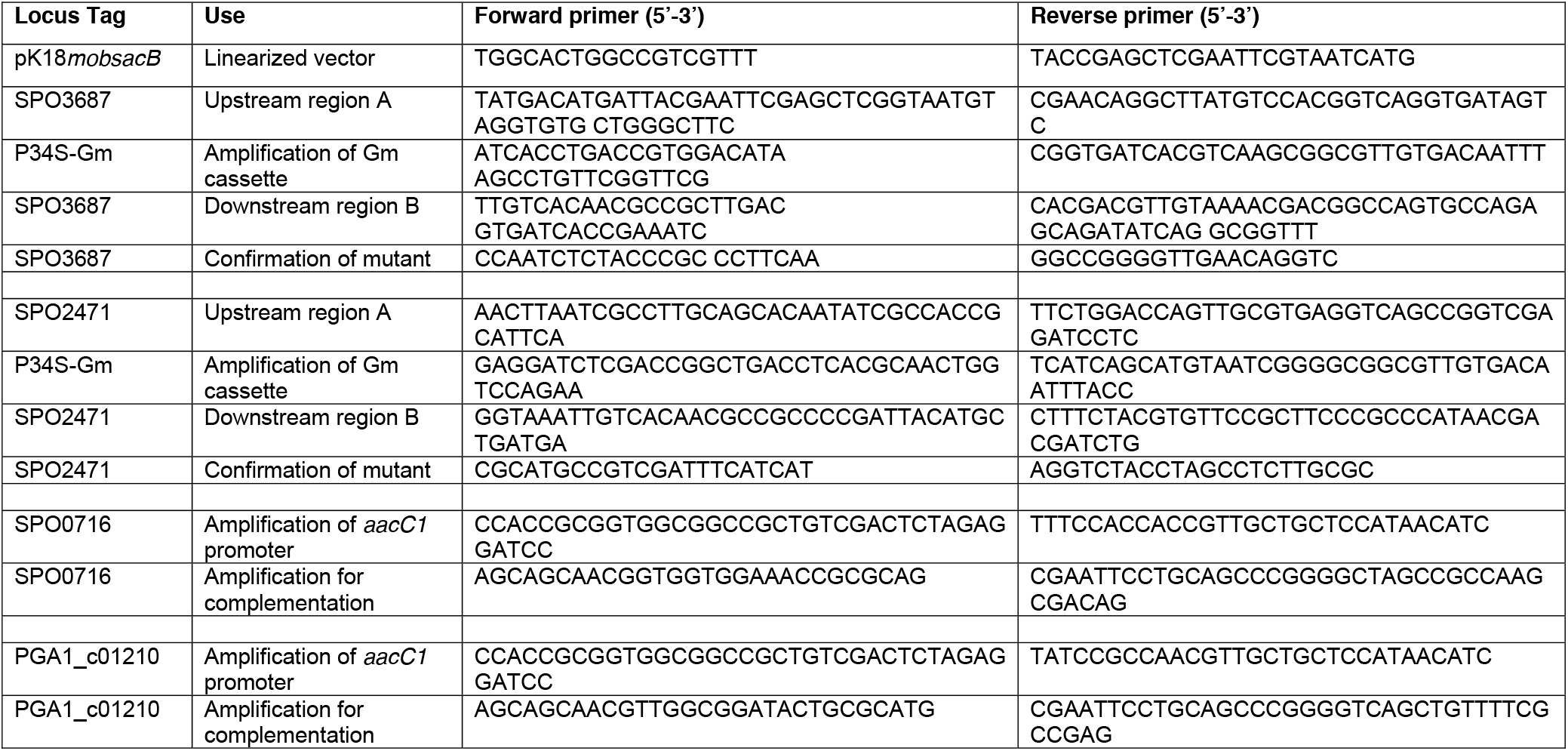
Primers used for molecular cloning and PCR amplification

**Figure S1.**
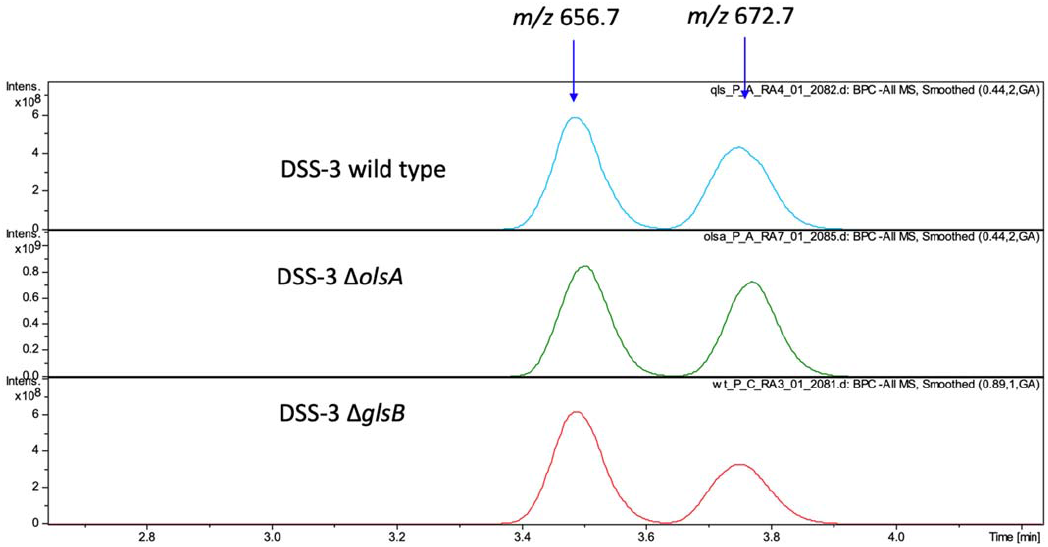
Sulfur-containing aminolipid (SAL) production in *Ruegeria pomeroyi* DSS-3 is not affected in the *olsA* mutant or the *glsB* mutant. The blue arrows point to the two major SAL species eluted at 3.4-4 min, with a *m/z* of 656.7 and 672.7 respectively. Cells were cultivated in ½ YTSS medium. Ions were collected in the negative (-ve) ionisation mode using electrospray ionisation (ESI) using an amaZon SL ion trap MS (Bruker)

**Figure S2.**
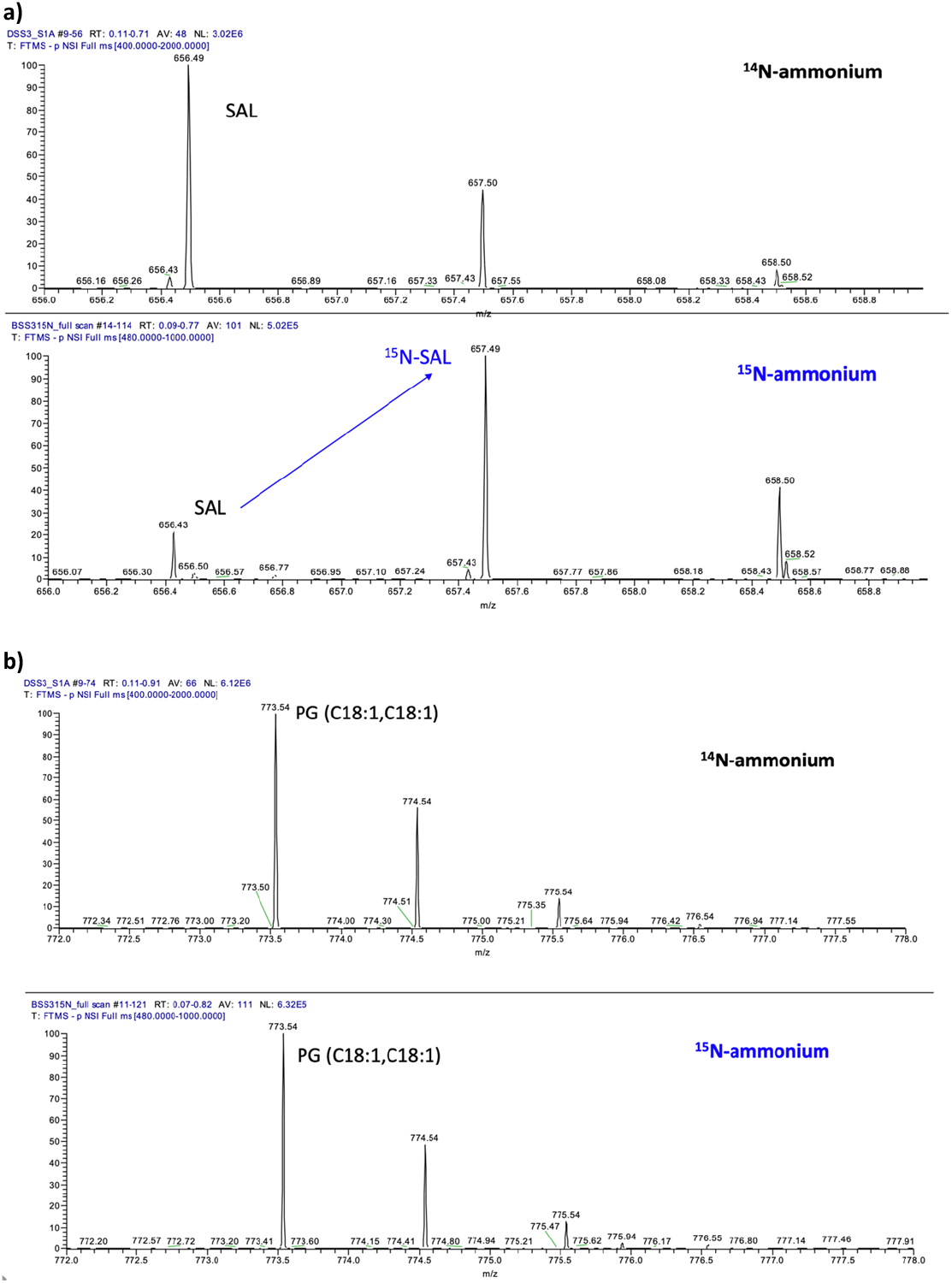
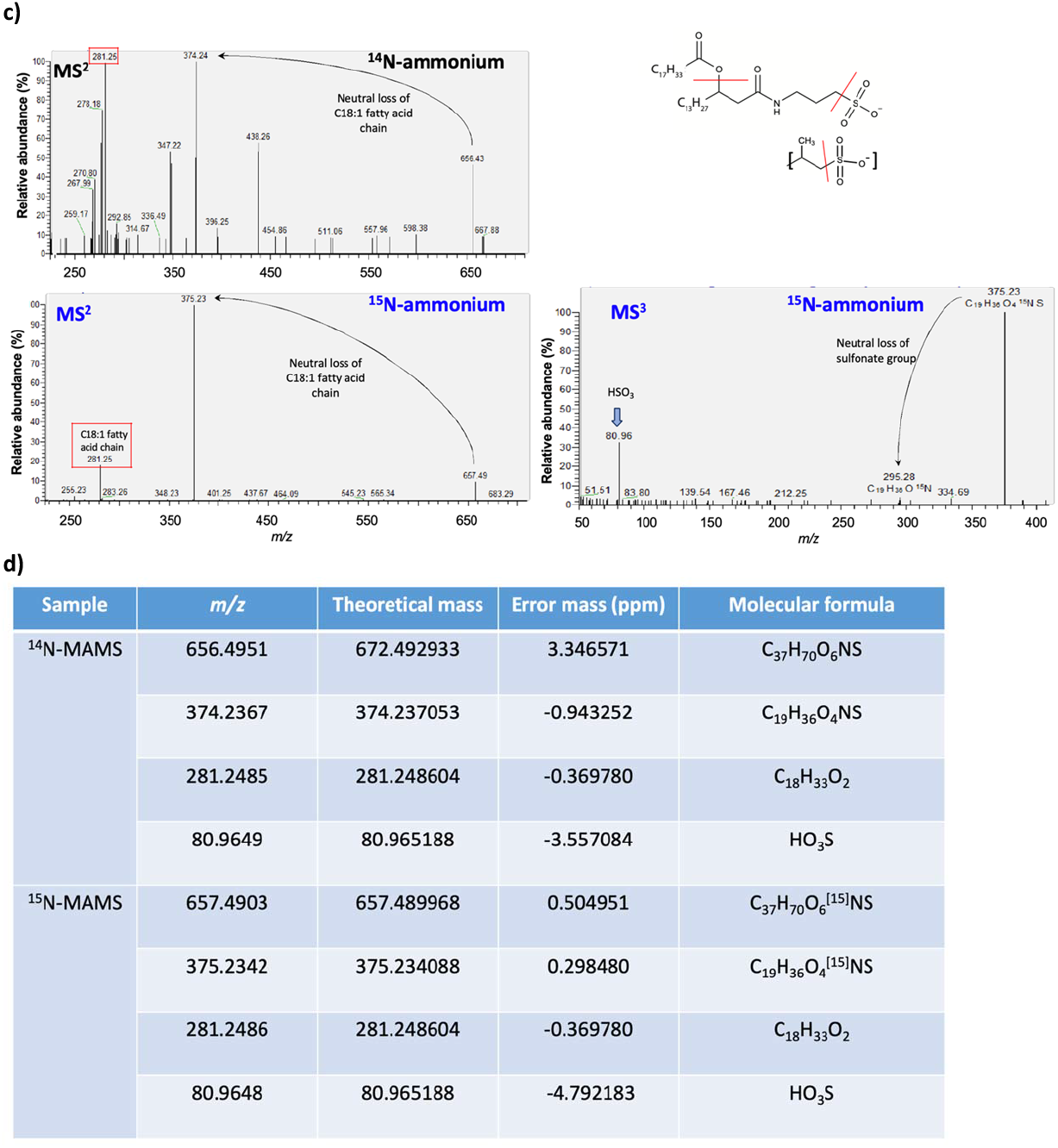
**a)** *Ruegeria pomeroyi* DSS-3 cells cultivated in the defined MAMS medium supplemented with ^15^N-labelled NH_4_Cl as the sole nitrogen source resulted in a significant enrichment of ^15^N-labelled SAL *(m/z* shift from 656.49 to 657.49) in the negative ionisation mode. **b)** However, no enrichment of ^15^N in the C18:1/C18:1 phospholipid PG was observed, which does not contain nitrogen. **c)** Further MS^n^ fragmentation of the ^15^N-labelled SAL *(m/z* 657.49) showed the production of a C18:1 fatty acid *(m/z* 281.25) and a *m/z* 375.23 ion which give rise to the formation of the HSO_3_ ion *(m/z* 80.96). Ions were collected and fragmented in negative (-ve) ionisation mode using an Orbitrap fusion MS (Thermo Fisher Scientific) by direct infusion. **d)** Theoretical mass and proposed molecular formula for the identified ions from *m/z* 656 SAL species.

**Figure S3.**
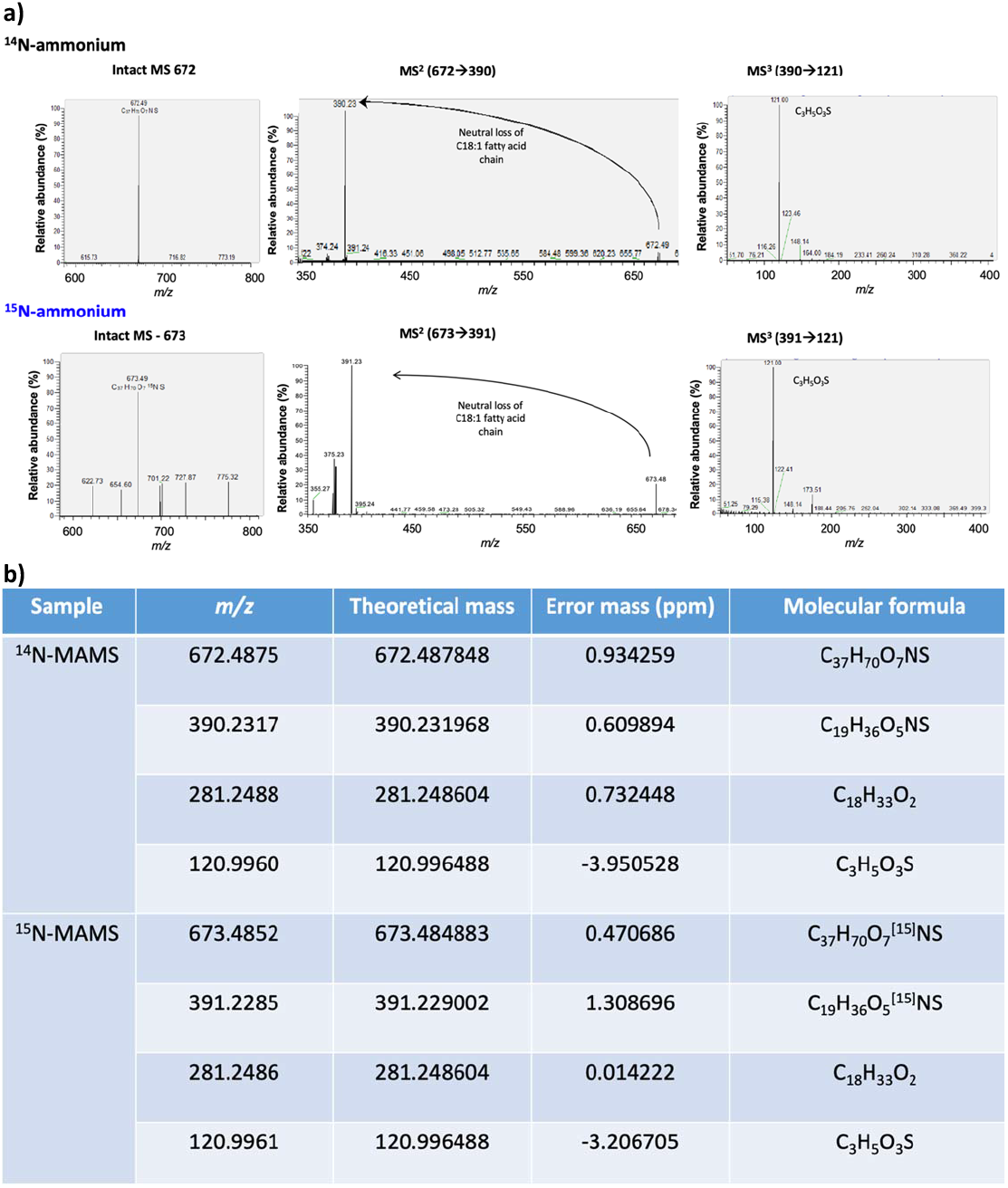
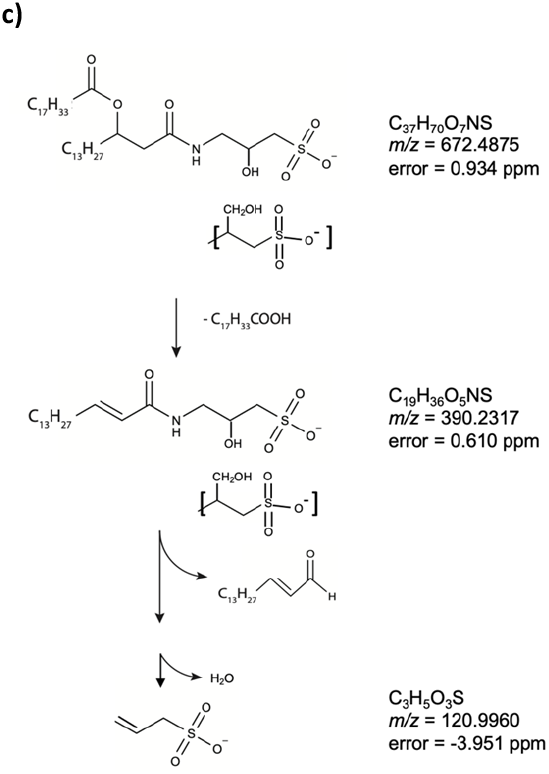
**a)** MS^n^ fragmentation of the *m/z* 672 and *m/z* 673 SAL lipid species extracted from *Ruegeria pomeroyi* DSS-3 cultivated in a defined MAMS medium supplemented with ^14^N-NH_4_Cl or ^15^N-NH_4_Cl as the sole nitrogen source, respectively. MS^n^ fragmentation was carried out in the negative ionisation mode using an Orbitrap fusion MS (Thermo Fisher Scientific) by direct infusion. **b)** Theoretical mass and proposed molecular formula for the identified ions are also shown. Both *m/z* 672 and *m/z* 673 SAL produced a sulfur-containing ion of *m/z* 121 (2-propene-1-sulfonate) and neither HSO_3_ ion *(m/z* 80.96) nor SO_3_ ion *(m/z* 79.96) was observed. **c)** Proposed fragmentation pattern of *m/z* 672 SAL species.

**Figure S4.**
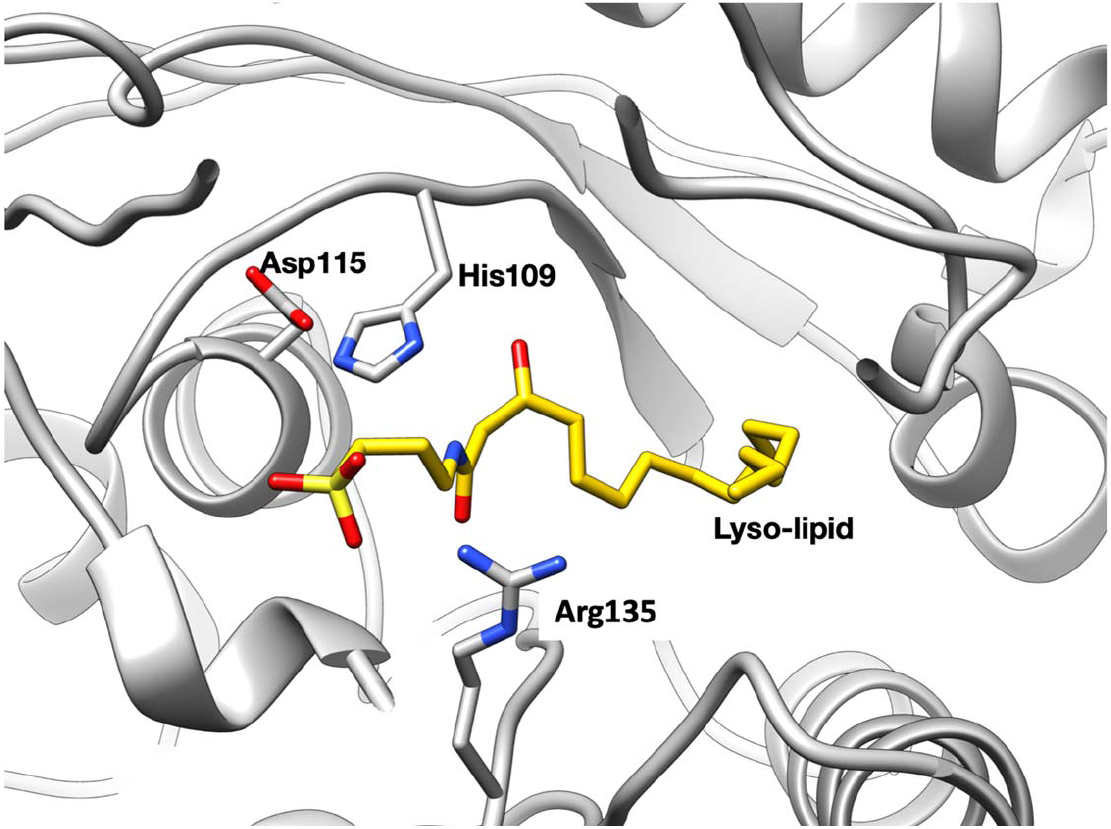
Homology model of SalA docked with the lyso-SAL lipid, showing a possible pose for the lyso-SAL lipid hydroxy group adjacent to His109 with the Arg135 suggested to coordinate the sulfonate head group.

**Figure S5.**
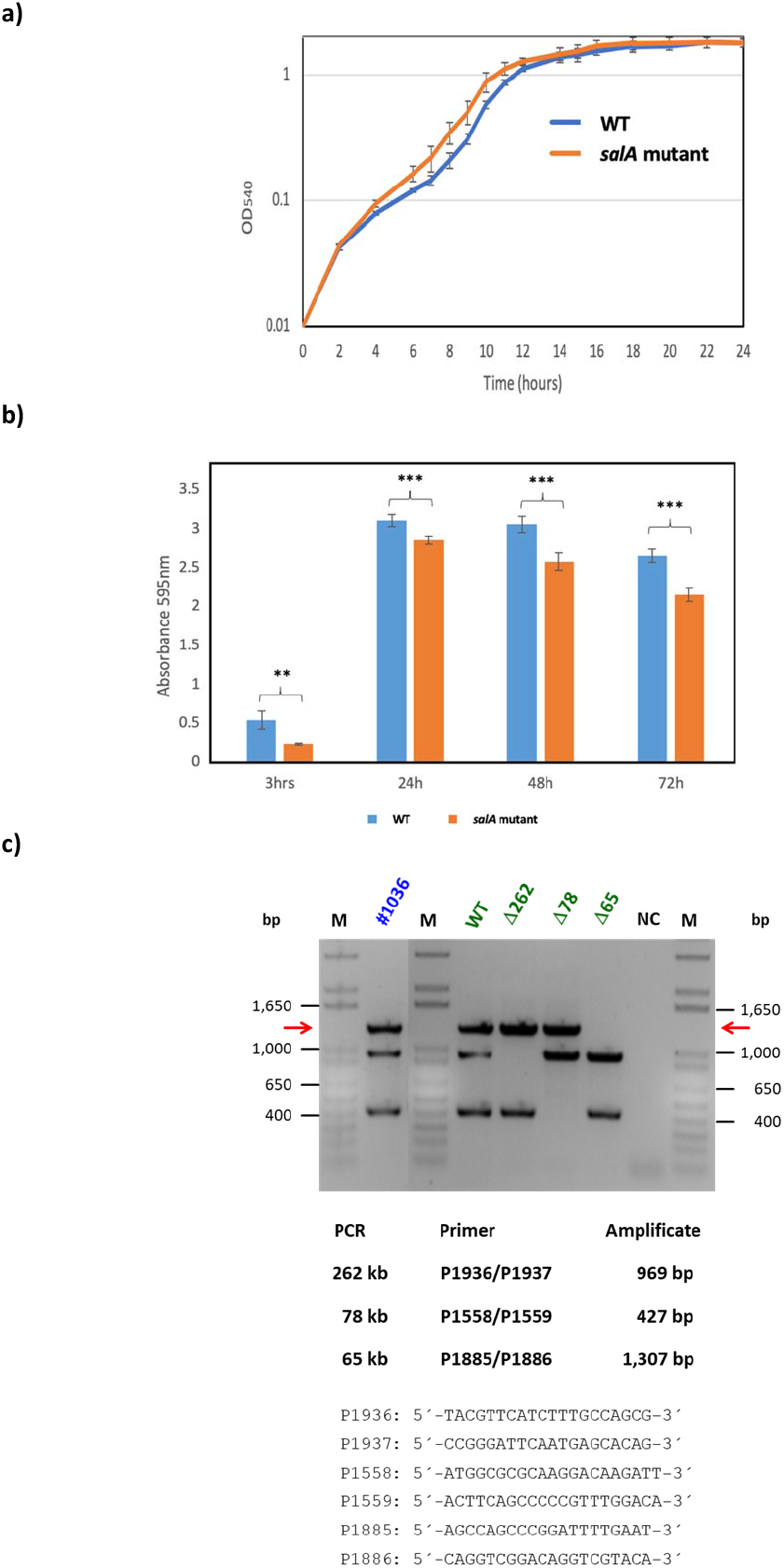
**a)** Growth of the wild-type and the *salA* mutant strain of *Phaeobacter inhibens* DSM 17395 in marine broth medium. **b)** Biofilm assay showing the differences in biofilm formation on plastic surfaces in marine broth medium between the wild-type and the *salA* mutant strain of *Phaeobacter inhibens* DSM 17395. The absorbance of crystal violet was measured at 595 nm after 3 h, 24 h, 48 h and 72 h post inoculation. **, *p*<0.01; ***, *p*<0.001. **c)** Plasmid profiling of the *salA* mutant [no #1036] from *Phaeobacter inhibens* DSM 17395. Triplex PCR for the presence of the 262 kb, 78 kb and 65 kb plasmids. The wild type (WT) and three plasmid curing mutants (Δ262, Δ78, Δ65) served as a reference. Red arrows indicate the 1,307 bp PCR product of the 65 kb biofilm plasmid. NC, negative control; M, marker (Gibco, 1 Kb Plus DNA Ladder).

**Figure S6.**
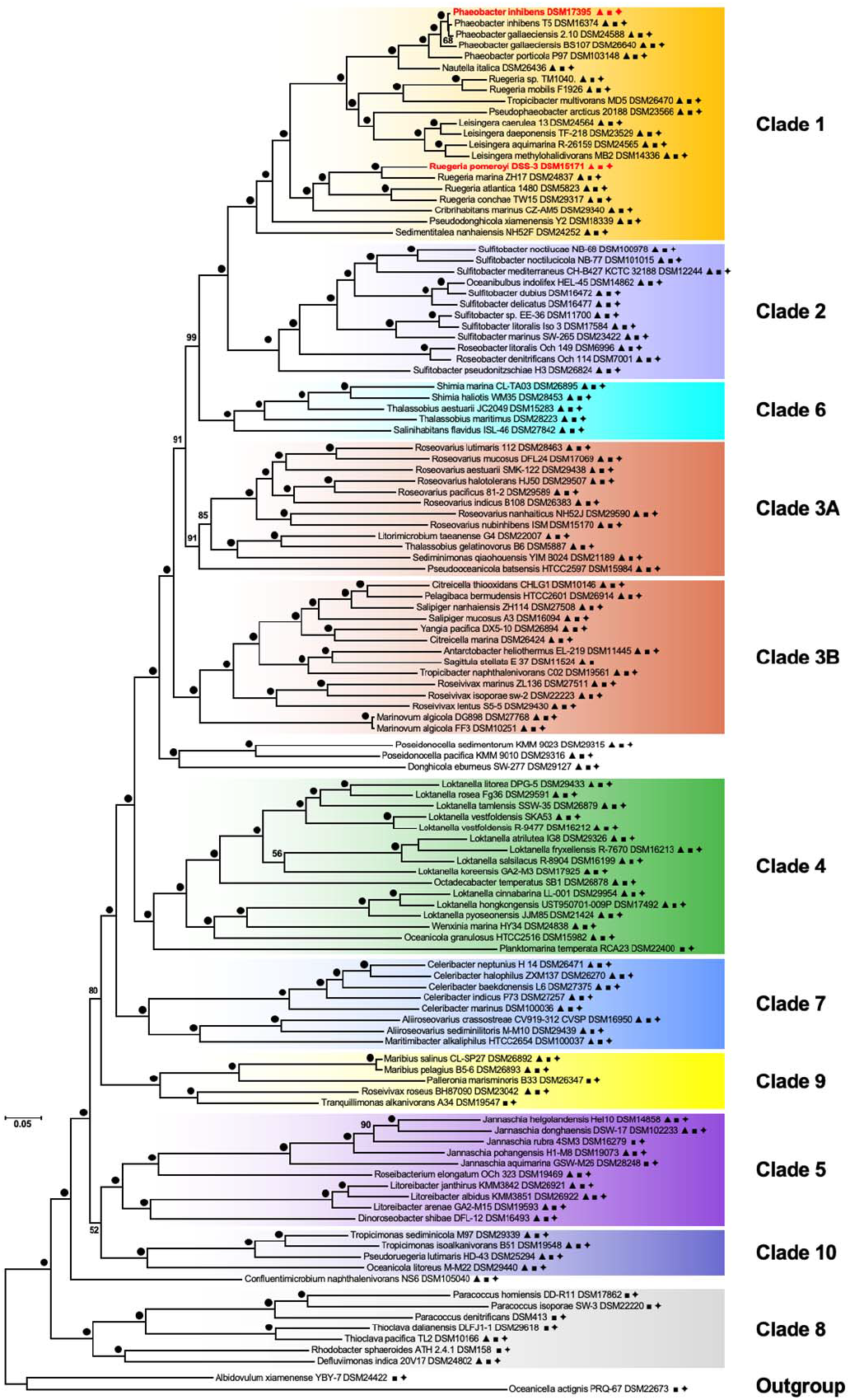
Distribution of *salA* (▲), *olsA* (■) and *pIsC* (✦) in genome sequenced roseobacters. The two model bacteria *Ruegeria pomeroyi* DSS-3 and *Phaeobacter inhibens* DSM17395 used in this study are highlighted in red. This phylogenomics tree was constructed by Bartling et al. (2018) using 120 genome-sequenced *Rhodobacteraceae* isolates with 504 universal marker gene alignments and a combined length of 153,625 conserved amino acid residues. Ten different lineages (Clade 1 to 10) were shown and the presence/absence of genes involved in sulfur-containing animolipid (*salA*), ornithine-lipid (*olsA*) and phosphatidic acid (*plsC*) synthesis is marked.

